# Cracking the case: Differential adaptations to hard biting dominate cranial shape in rat-kangaroos (Potoroidae: *Bettongia*) with divergent diets

**DOI:** 10.1101/2025.08.18.670733

**Authors:** Maddison C Randall, Vera Weisbecker, Meg Martin, Kenny Travouillon, Jake Newman-Martin, D. Rex Mitchell

## Abstract

Functional adaptation in the mammalian jaw is often best predicted by the hardest bites an animal makes. Therefore, even when closely related species have otherwise divergent diets, a shared biomechanically challenging resource should be reflected in similar adaptations to jaw biomechanics. We assessed this in two species of rat-kangaroos, whose otherwise divergent diets include the extremely tough-shelled seeds of *Santalum* spp. (sandalwood and/or quandong). We used geometric morphometrics to analyse cranial shape of 161 bettongs (*Bettongia* spp.), including all four extant species. We identified adaptations to higher bite forces in both species that crack open *Santalum* seeds. However, *B. lesueur* had shorter facial proportions, indicating higher mechanical advantage, while *B. penicillata* had a premolar morphology that likely focussed bites to a specific, reinforced position on the jaw. This represents an example of many-to-one mapping at the genus level. We also found differences in a subsample of captive northern bettongs compared to wild conspecifics, suggesting some role of phenotypic plasticity in shaping adult skulls. The large olfactory tracts of *B. penicillata*, that support search for underground fungi, might have constrained its cranium to retain longer proportions. Fungal abundance could potentially be an important consideration in identifying translocation sites for this species.

## INTRODUCTION

In the study of vertebrate skulls, it is becoming increasingly clear that skull morphology is better predicted by performance extremes than the commonly applied, broad diet classifications (Figueirido, Tseng & Martin-Serra, 2013; Mitchell, 2019; Santana, Dumont & Davis, 2010; Zelditch, Ye, Mitchell & Swiderski, 2017). This is probably because bone structure must include safety factors to withstand these extremes (Alexander, 1981; Willie, Zimmermann, Vitienes, Main & Komarova, 2020). Blanket diet categories (such as “grazer”, or “fungivore”) are often too vague to acknowledge small proportions or seasonal reliance on crucial food groups that might often be more challenging to obtain or process than the overall category would suggest (e.g., ‘fallback foods’) (Constantino & Wright, 2009). On top of cyclical strains from frequent, repetitive biting behaviours (Hylander & Johnson, 1997), peak strains from more demanding behaviours are also expected to influence bone deposition across micro- and macroevolutionary scales (Biknevicius & Ruff, 1992; Frost, 1994). The evolution of a species’ skull morphology should therefore also be influenced by their most biomechanically challenging foods or biting behaviours used to access resources (whichever places more strain on the skull), even if they are eaten rarely or in small quantities.

An influence of biomechanically taxing foods on cranial adaptation might result in the evolution of morphologically convergent cranial architectures (i.e., similarities that are not found in the common ancestor of the species (Stayton, 2015)) in species that otherwise consume very different diets. This is conceptually aligned with Liem’s Paradox, which states that species that appear specialised to particular foods can frequently have generalist diets (Robinson & Wilson, 1998). In such cases, morphology can reflect the demands of less-preferred foods that are vital for overall survival but might only be selected in the absence of more preferred resources (Robinson & Wilson, 1998; Ungar, Grine & Teaford, 2008). It therefore potentially represents a crucial piece in understanding processes of functional convergence.

Small rat-kangaroos of the genus *Bettongia* are an excellent test case for this ‘biomechanical extremes’ hypothesis. Bite simulations done on the genus using finite element analysis (FEA) have shown high diversity in mechanical advantage and stress magnitudes, suggesting that skull shape might be related to the variable demands of their diets (Mitchell, Wroe, Martin & Weisbecker, 2025). All members of the genus are specialists at locating and consuming low fibre, high nutrient foods. This includes food groups with considerably different material composition, such as fruits, fungi, seeds, roots, tubers, plant matter, and insects (Arman & Prideaux, 2015; Bice & Moseby, 2008; McIlwee & Johnson, 2002; Mitchell, Ledogar, Andrew, Mathewson, Weisbecker & Vernes, 2024a; Robley, Short & Bradley, 2008; Taylor, 1992; Vernes, Castellano & Johnson, 2002; Zosky, Wayne, Bryant, Calver & Scarff, 2017).

However, only two species, the burrowing bettong (*B. lesueur*) and the brushtail bettong (*B. penicillata*), crack open extremely tough-shelled *Santalum* seeds (Chapman, 2015; McNamara, 2014; Murphy, Howard, Hardy & Dell, 2015; Murphy, Garkaklis & Hardy, 2005). *Bettongia lesueur* lives in Australia’s arid zones, and *B. penicillata* lives in temperate and semi-arid regions, (Fig. 1A) but they are not sister taxa (Fig. 1B). The remaining two living species, the Tasmanian bettong (*B. gaimardi*) and the northern bettong (*B. tropica*) are located in more temperate or tropical regions of Australia (Fig. 1A) and are not known to consume *Santalum* seeds, as their distribution does not overlap with the species.

**Figure 1:**
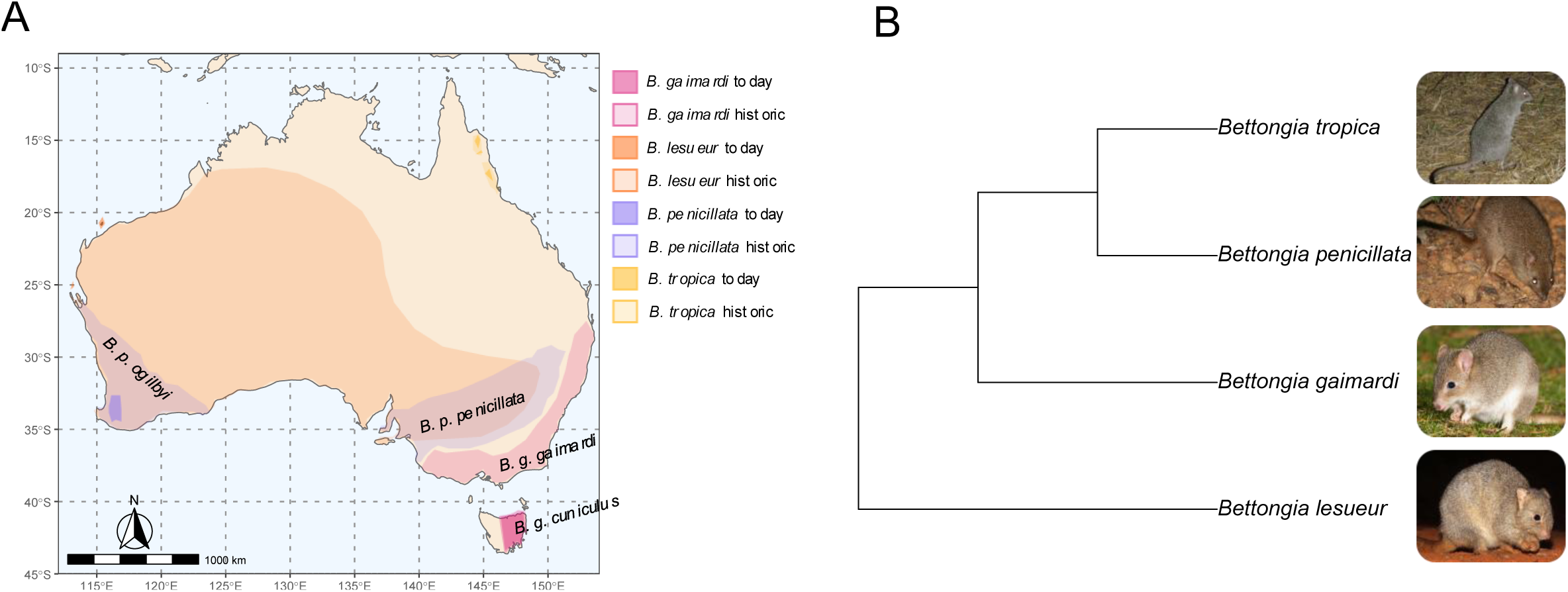
(A) Current and historic distributions of extant *Bettongia* spp. (Haouchar et al., 2016; Rick, Ottewell, Lohr, Thavornkanlapachai, Byrne & Kennington, 2019) (B) Phylogeny of all extant *Bettongia* spp. (Westerman, Loke, Tan & Kear, 2022). Photo credit: *B. penicillata*, Authur Chapman CC BY 4.0; via Flickr. *B. lesueur*, Tom Hunt CC BY NC 4.0; via Lex. *B. tropica*, “Kaitlyn”, and *B. gaimardi*, J.J Harrison CC BY 4.0; via Wikimedia Commons

Aside from the consumption of *Santalum* seeds, the diet composition of the brushtail and burrowing bettongs differs substantially (Haouchar, Pacioni, Haile, McDowell, Baynes, Phillips, Austin, Pope & Bunce, 2016; McDowell, Haouchar, Aplin, Bunce, Baynes & Prideaux, 2015). The burrowing bettong has a variable diet with greater reliance on more mechanically resistant roots, seeds, and browse vegetation (Arman & Prideaux, 2015; Mitchell et al., 2024a; Robley et al., 2008) than the brushtail bettong, which has a greater dependence on softer fungal sporocarps (i.e., truffles) (Mitchell et al., 2024a). However, their shared consumption of *Santalum* seeds means that both species are adapted to some of the most difficult food items to bite onto. The outer casings (testae) of *Santalum* seeds often reach 20mm in diameter and have mechanical properties of toughness and stiffness that exceed those of other known resistant food materials, such as cherry pits and popcorn kernels (Mitchell et al., 2024a); in fact, a quandong (*Santalum acuminatum*) seed case required at least 100kg of direct force to crack using material testing equipment (Mitchell et al., 2024a).

At only around two kg of body mass (Page, Ruykys, Miller, Adams, Bateman & Fleming, 2019; Rose & Rose, 1998), and skulls typically measuring a maximum length of ∼70mm (McDowell et al., 2015; Rose & Rose, 1998), the capability of these rat-kangaroos to extract the nutritious kernels from *Santalum* casings is remarkable. The relatively large, sectorial premolars of potoroids are the main site of mechanical pre-processing of most foods (McNamara, 2014; Sanson, 1989). A large, elongate third premolar can be traced back in the fossil record to the Oligocene and Miocene, when woodlands and rainforests were more widespread in Australia (Jackson et al. 2024). Similar premolars are found in ancient macropodids genera from the Oligocene and Miocene specifically in *Bulungamaya* and *Gumardee* (Flannery, Archer & Plane, 1983); *Cookeroo* (Butler, Travouillon, Price, Archer & Hand, 2016); *Ganguroo* (Cooke, 1997); *Ngamaroo* (Kear & Pledge, 2007); *Purtia* (Case, 1984); *Wabularoo* (Archer, 1979); and *Wakiewakie* (Woodburne, 1984). In Potoroidae, it is found in the middle Miocene *Bettongia moyesi* (Flannery & Archer, 1987) and the upper Pleistocene *Borungaboodie hatcheri* (Prideaux, 1999). This highlights how this trait is an ancestral adaptation that is retained in the modern bettongs, and it is also the location of the tooth row where the bettongs crack open *Santalum* seeds. This involves repetitive hard biting onto the seed with premolars, while rotating the seed with the forelimbs between bites. The process has been detailed for *B. penicillata* and appears to involve considerable effort and extensive mouthing of the seed, as if propagating and monitoring microfractures in the casing (McNamara, 2014).

Despite the shared consumption of *Santalum* seeds in burrowing and brushtail bettongs, previous works based on linear measurements have noted differences between bettong species in the skull which are thought to be the result of variation in habitat and dietary preferences (Haouchar et al., 2016; McDowell et al., 2015). However, dietary specialisations are known to also drive convergent evolution of the skull (e.g., Figueirido, Serrano-Alarcón, Palmqvist & Kitchener, 2011; Klaczko, Sherratt & Setz, 2016; Samuels, 2009; Santana, Grosse & Dumont, 2012), with adaptations to generating higher bite forces tending to be more consistent across taxa (Figueirido et al., 2013; Sansalone, Wroe, Coates, Attard & Fruciano, 2024). We might therefore expect some similarity between brushtail and burrowing bettongs if the extreme efforts required to crack *Santalum* seeds are reflected in evolution of their skull. Thus, convergence in skull morphology would exhibit features indicative of higher bite forces, such as a shorter rostrum or more robust features, compared to bettongs that consume softer diets (Mitchell et al., 2018; 2024).

In this study, we test the above expectations using geometric morphometrics to analyse cranial shape of all four species of extant bettongs, the *Santalum*-seed-cracking burrowing bettong and brush-tail bettong and the Tasmanian and Northern bettong that do not crack these seeds. We expected to find structural adaptation to generating and supporting high bite forces in the two species known to crack *Santalum* seeds. We therefore expected to find either similar morphologies or biomechanically equivalent adaptations to hard biting.

## MATERIAL AND METHODS

Our sample included the crania from 161 adult specimens from all four extant species of *Bettongia*: *B. gaimardi* (n=33), *B. lesueur* (n=39), *B. penicillata* (n=77), *B. tropica* (n=12). We identified adults through eruption of the P3 and M4 (Janis, 1990; Mein, Manne, Veth & Weisbecker, 2022; Mitchell, Sherratt, Ledogar & Wroe, 2018). 35 of the 78 *B. penicillata* were of the subspecies *B. p. ogilbyi*, originating from the western historic distribution of the species (Fig. 1A). However, the remainder of the sample for this species were a combination of reintroduced *B. p. ogilbyi* and native *B. p. penicillata* from more eastern localities – the majority of which lacked taxonomic distinctions to a subspecies level. The sample of wild *B. tropica* crania was small (n=3), but we also included crania from some captive-bred individuals (n=8) of this endangered species. Additionally, *B. penicillata* specimen M18184 was collected from Queensland in the 1860’s and is therefore almost certainly *B. tropica*. We allocated it as such, as a 12^th^ specimen. We sourced specimens from the Western Australian Museum, South Australian Museum, Tasmanian Museum and Art Gallery, Queen Victoria Museum, Australian Museum, Australian National Wildlife Collection & and Queensland Museum.

We generated 3D surface models of each cranium *in silico* (Viacava, Blomberg, Sansalone, Phillips, Guillerme, Cameron, Wilson & Weisbecker, 2020). Most specimens were 3D scanned using a Polyga compact S1 scanner (Polyga, Vancouver, Canada), however, we also scanned two specimens of *B. tropica* using a Nikon XT H 225ST CT Scanner (Nikon Metrology, Tring, UK). Nine specimens had unilateral damage to the zygomatic arch, making landmark placement impossible from those locations. However, including damaged specimens is unlikely to appreciably impair analyses of gross morphology and dominant predictors of shape (Mitchell, Kirchhoff, Cooke & Terhune, 2021). Thus, we repaired these specimens by superimposing mirrored sections from the undamaged side using Geomagic Wrap (Table S1; 3D Systems, Rock Hill, South Carolina).

Each 3D model was landmarked by lead author MR with Checkpoint (Stratovan, Sacramento, CA, USA) using a predefined template (Mitchell, Potter, Eldridge, Martin & Weisbecker, 2024b; Viacava et al., 2020) (Table S2) comprising 147 landmarks (99 fixed landmarks, and four curves of semi-sliding landmarks). This landmarking protocol is adapted from Viacava et al., (2020; 2022) to capture functionally relevent aspects of the cranial morphology influenced by dietary pressures. For example our landmarks capture relative rostum length and skull width which are known to respond to dietary pressure. To ensure the repeatability of the manually placed landmarks, fifteen specimens were digitised three times. Contributing a total of 5.6% of variation, measurement error was much lower than interindividual variation.

All analyses were done in R statistical software (R Development Core Team, 2023), using the Morpho (Schlager, Jefferis & Ian, 2018) and geomorph (Adams, Otárola-Castillo & Paradis, 2013) R packages. We initially replaced any missing landmarks using the ‘estimate.missing’ function from geomorph. We then used a generalised Procrustes analysis (GPA) to remove the degrees of freedom associated with size, rotation, and translation (Rohlf & Slice, 1990).

We then removed shape variation explained by asymmetry (Klingenberg, Barluenga & Meyer, 2002; Mardia, Bookstein & Moreton, 2000). The symmetrical component of shape was used in all further analyses. For linear models, we used Procrustes regression modelling and Procrustes ANOVAs, implemented by geomorph. This uses a random permutation approach to assess the significance of the model (Collyer & Adams, 2018).

Assessment of interspecific differences in cranial shape requires an investigation of whether observed differences might be explained by differences in size and size-correlated changes of shape (allometry) (Klingenberg, 2016). We therefore assessed Procrustes regression of cranial shape and log-transformed cranial size (centroid size) to identify the overall contribution of allometry in the sample relative to species differences. We also considered potential interaction effects of size and species (shape ∼ log(size)*species) that would indicate interspecific differences in allometric slopes.

We also used Principal Components Analysis to see if species differentiation was the main driver of variation in our sample, with the expectation that the main axes would reflect differences between species. We performed regressions between PC scores and log-transformed cranial centroid size to exclude a common allometric effect as the main source of interspecific variation. To visualise shape changes reflected in the first two principal components, we warped a mesh of the specimen closest to the sample consensus shape to the shape predicted by the extremes of each principal component to accompany the PC plots.

To best visualise interspecific shape differences with allometric effects accounted for, we first tested whether species means of cranial centroid size differed across species (log(size) ∼ species). If there was no significant difference in mean centroid size, we could warp a surface mesh of each species to the mean shape of their respective species. However, if mean centroid size differed significantly, we would warp species meshes to cranial shapes predicted by an identical value of centroid size, to remove as much allometry as possible in interspecific visualisations.

## RESULTS

Our Procrustes regression shows that species differences account for most of the shape variation in the sample (37.2%), followed by allometric effects (13.3%) (Table 1). In addition, there is a significant interaction effect of species and size, indicating some differences in allometric slopes between species. Interspecific differences in allometric slopes can be seen in Figure 2A, with the slope of *B. penicillata*, possibly indicating a faster rate of allometric change across its range of cranial size than the other species or perhaps representing more than one taxon. However, the interaction effect explains a very small amount of the variation (1.5%), so we deemed it acceptable to compare species without expecting a confounding effect of the interaction in our PCA analyses. Based on our regression, we expected species differentiation to play a major role and allometry a subordinate role. Pairwise comparisons of species show that all species are significantly different in shape, both in a univariate model and when factoring in cranial size (Table 2).

**Table 1:** Results of our Procrustes ANOVA of shape associations with size, species, and interaction term between size and species.

| **Response** | **Predictor** | **df** | **R<sup>2</sup>** | **F** | **p** |
| --- | --- | --- | --- | --- | --- |
| Shape | log(cranial centroid size) | 1,153 | 0.133 | 42.205 | **0.001** |
|  | species | 3,153 | 0.372 | 39.436 | **0.001** |
|  | log(cranial centroid size) x species | 3,153 | 0.015 | 1.542 | **0.002** |

**Table 2:** P-values of interspecific differences in shape. All comparisons are significant (*p* ≤ 0.001) both alone, and when factoring in cranial size.

| **shape ~ species** |  |  |  |
| --- | --- | --- | --- |
|  | *B. gaimardi* | *B. lesueur* | *B. penicillata* |
| *B. lesueur* | **0.001** |  |  |
| *B. penicillata* | **0.001** | **0.001** |  |
| *B. tropica* | **0.001** | **0.001** | **0.001** |
| **shape ~ log(size) + species** |  |  |  |
|  | *B. gaimardi* | *B. lesueur* | *B. penicillata* |
| *B. lesueur* | **0.001** |  |  |
| *B. penicillata* | **0.001** | **0.001** |  |
| *B. tropica* | **0.001** | **0.001** | **0.001** |

**Figure 2:**
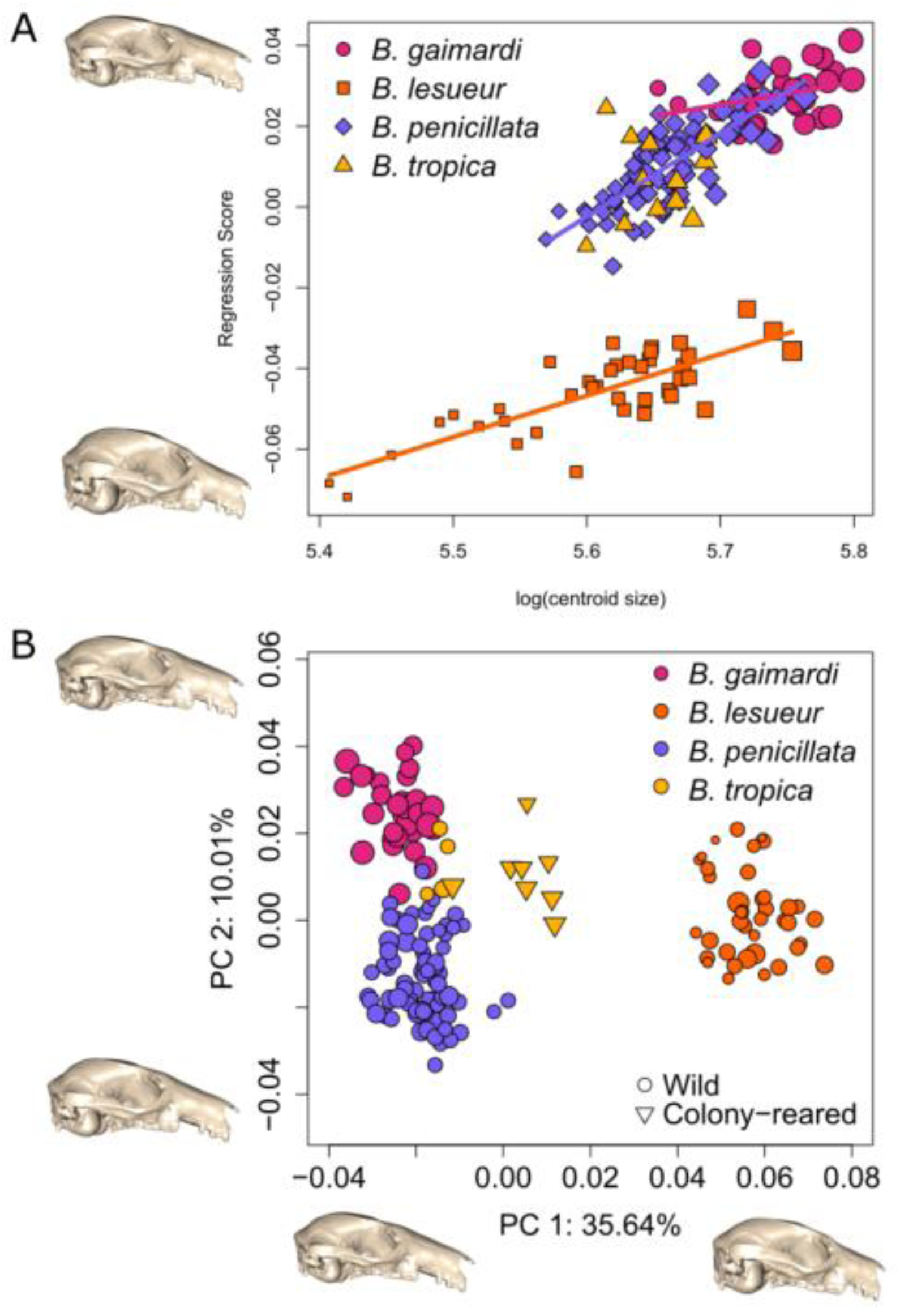
(A) Interspecific allometric gradients *(B. tropica* excluded from analysis due to small sample size and narrow size range) (B) Principal component analysis (PCA) shows where dominant shape differences are that distinguish species (point size represents cranial size). The captive-reared *B. tropica* are noticeably isolated from the wild *B. tropica* that cluster closer to *B. gaimardi*.

**Figure 3:**
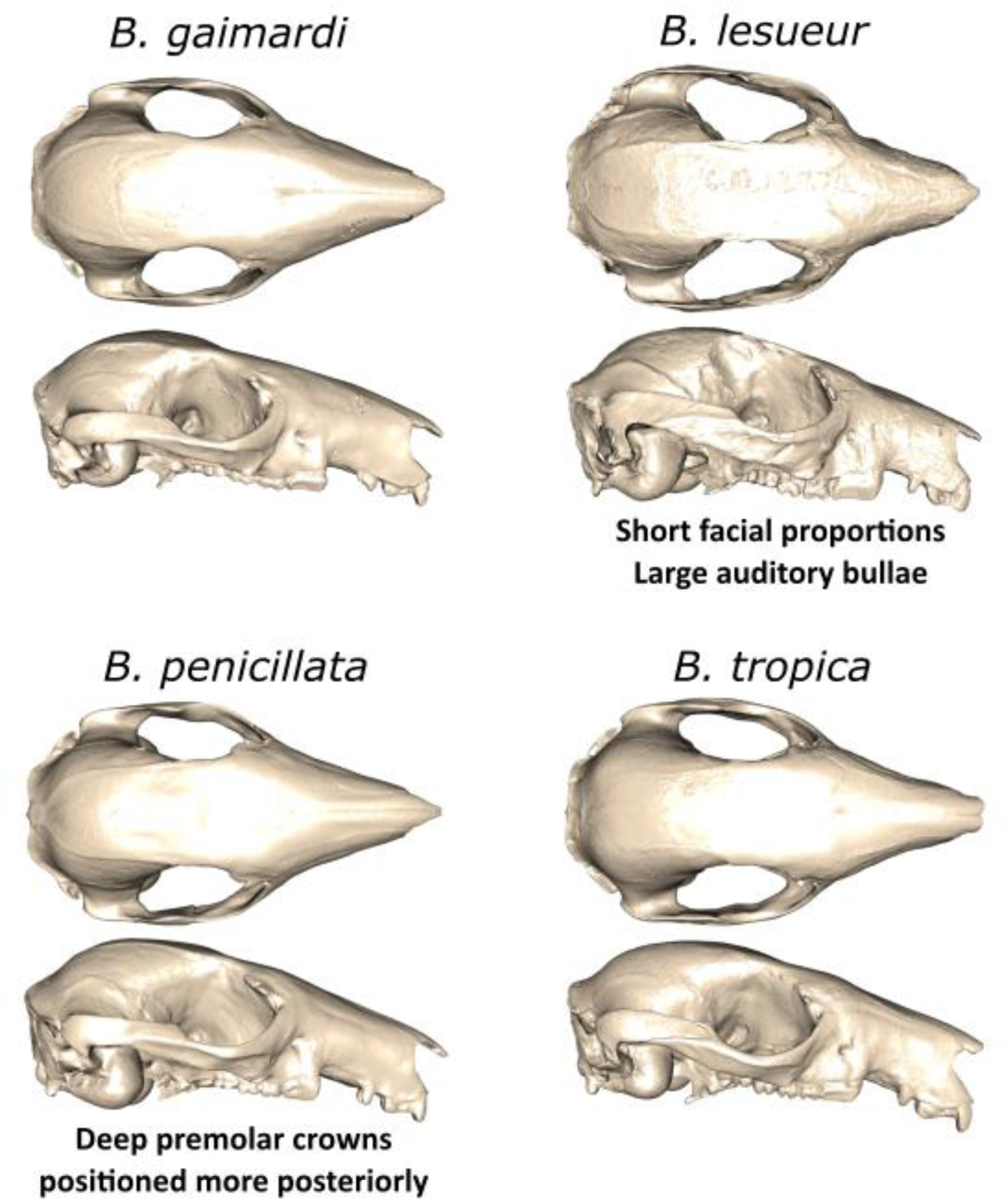
Predicted shape for all four species at identical cranial size (determined from centroid sizes on allometric gradients). PC1 of Figure 2B identifies the mechanically efficient facial proportions of *B. lesueur*, while PC2 identifies the shape and position of the premolars in *B. penicillata*.

A principal component analysis of cranial shape variation (Figure 2B) shows that the main variation is dominated by differences between species. There is no visible further separation between species on other PCs.

PC1, representing 35.64% of all shape variation, separated *B. lesueur* from all remaining extant *Bettongia* spp. The maximum scores of PC1, occupied solely by *B. lesueur*, indicated that this species differs from all others in having a shorter rostrum, more robust zygomatic arches, and enlarged auditory bullae (Fig. 2B). PC1 correlated significantly with cranial size (R^2^=0.243, p=0.001), but adjusting for allometry is not useful in this case because of the substantial intercept difference between *B. lesueur* and the remainder of the sample. The allometry in PC1 is likely associated with the range of smaller cranial sizes for *B. lesueur* (Fig. 2A), but there is no indication that this effect confounds the overall clear differentiation between species.

PC2, representing 10.01% of total shape variation, largely separated *B. penicillata* from *B. gaimardi* and *B. tropica*. This component mostly described the anteroposterior length and positioning of the P3. The minimum of PC2, represented by *B. penicillata*, described premolar crowns with a slicing edge reduced in length that were also positioned more posteriorly in the oral cavity. The maximum region of the PC2 space was occupied by *B. gaimardi*, while the *B. tropica* are clustered towards the middle of this space. PC2 also correlated with cranial size (R^2^=0.053, p=0.003). However, even though the maximum of PC2 was occupied by the cluster of species with the largest crania, *B. gaimardi*, the smallest individuals in the sample, members of *B. lesueur* (Fig. 2A), did not represent the minimum extreme of PC2 (Fig. 2B), suggesting that variation on PC2 is more attributable to other factors.

The interspecific test of mean centroid sizes showed that species differed significantly in centroid sizes (R^2^=0.459, p=0.001). As described in the methods, to visualise cranial shape across species, with minimal effects of allometric differences, we therefore scaled meshes of each species to their respective cranial shapes predicted for a centroid size shared by all species - a log(size) of 5.68 (Fig. 2). The shape differences defined by the first two principal components, the shorter faces/larger bullae of *B. lesueur* and more posterior premolars of *B. penicillata*, are clearly visible in the warps.

## DISCUSSION

As expected, the two species of bettongs known to crack open the mechanically challenging outer casings of *Santalum* seeds, *B. lesueur* and *B. penicillata*, have different morphology of the masticatory apparatus compared to the two species that do not crack open these seeds due to no overlap in distribution with *S. acuminatum* or *S*. *spicatum*. However, counter to our expectations, the two *Santalum*-cracking species did not have similar adaptations to generating and withstanding high bite forces. This points to the independent acquisition of seed predation and associated adaptations in *B. lesueur* and *penicillata*. The evolutionary origin of this convergence depends on the ancestral condition, which we could not ascertain due to a lack of closely related relatives. However, seed predation and associated adaptations must have originated independently, either in both groups (if the ancestral shape was like *B. gaimardi/tropica*) or, either in both groups (if the ancestral shape was like *B. gaimardi/tropica*) or *Santalum*-cracking behaviours were secondarily regained in *B. penicillata* (if the ancestral condition was like *B. lesueur*).

Our results align with Liem’s paradox, which posits that mechanically extreme dietary items can drive skull shape evolution and obscure a much less strenuous average diet. Even if consumed infrequently, mechanically highly demanding foods like Sandalwood seeds are likely to place the greatest mechanical stress on the skull, potentially eliciting both short-term and long-term impacts on skull morphology (Robinson & Wilson, 1998; Ungar, Grine & Teaford, 2008). Inability to handle high biting forces – leading to skull, jaw or tooth fracture – could be fatal, thus favouring the evolution of a more robust cranium. This suggests a trade-off between the energetic costs of maintaining robust biological structures and the survival advantage of being able to process extreme, but rarer food items (Mitchell et al., 2024a).

The genetic and dietary divergence of *B. lesueur* (Haouchar et al., 2016; Mitchell et al., 2024a) is clearly reflected in shape differences between this species and the other three. Aside from larger auditory bullae, the main differences related to the short, robust proportions of its facial skeleton. This is consistent with findings from FEA that *B. lesueur* has the highest mechanical advantage of the genus (Mitchell et al., 2025). Shorter faces are frequently associated with high mechanical advantage, which permits conversion of more muscle force to bite force (Goswami, Milne & Wroe, 2010; Mitchell, 2019; Mitchell et al., 2018; Wroe & Milne, 2007). The deeper zygomatic arches in *B. lesueur* might represent a higher capacity to withstand forces exerted by the masseter complex during hard biting (Mitchell, 2019). This morphology would be expected in species capable of biting extremely hard, tough, or otherwise strong materials.

The remaining three species all differed from *B. lesueur* in having overall longer faces, despite only *B. penicillata* cracking *Santalum* seeds. The longer face of *B. penicillata* translates into a lower mechanical advantage; however, recent FEA revealed similarly low strain in this species to *B. lesueur* during simulations of equivalent bites (Mitchell et al., 2025). This result could be explained by a greater degree of bone deposition either as part of its developmental programme (Frost 1994, Pearson & Lieberman 2004, Ruff et al. 2006) or phenotypic plasticity (Weisbecker et al. 2019, Mitchell et al. 2020, Mitchell et al. 2021b).

However, *B. penicillata* additionally differed in shape from *B. gaimardi* and *B. tropica* at the upper third premolar (P3), the site where cracking *Santalum* seeds open takes place. The premolar crowns of *B. penicillata* are shorter in anteroposterior length, and deeper, creating a more chisel-like appearance compared to the more blade-like shape seen the other three *Bettongia* spp. The P3 is also positioned more posteriorly in the oral cavity in *B. penicillata*. Therefore, although the longer face of *B. penicillata* has a lower mechanical advantage than *B. lesueur*, its premolar morphology focusses the bites at these teeth to a smaller area with deeper crowns.

For both *B. lesueur* and *B. penicillata*, it is difficult to determine the degree to which selection for *Santalum*-cracking abilities has shaped the evolution of the cranium. For example, the robust shape of the *B. lesueur* cranium is probably not solely the result of adaptations for *Santalum* cracking. Rather, it might relate to the species’ overall greater dependence on more mechanically rigorous roots, seeds, and browse vegetation (Bice & Moseby, 2008; Mitchell et al., 2024a; Robley et al., 2008). By contrast, the clear difference in premolar shape in *B. penicillata* suggests a substantial contribution of evolved adaptation to hard cracking in the context of selection for an otherwise longer face. Defining clear relationships between morphology and behaviour is further complicated by the fact that the way by which bettongs can crack open *Santalum* seeds has only been formally described for *B. penicillata* (McNamara, 2014). This species is described as relying heavily on the third premolars and applying substantial effort to access the kernels inside the tough testae. While it is likely that *B. lesueur* applies a similar approach, we cannot yet rule out other possible behaviours involving its greater mechanical advantage with the front teeth. Mechanical analyses of the material properties additionally found that the testae are stiffer and less tough after drying, which would be easier to crack open. This might explain the scatter-hoarding or “caching” seen in both species (Mitchell et al., 2024a).

An additional challenge to ecological interpretation of mammalian crania arises from the fact that our species display functional convergence without showing associated morphological similarities. In particular, facial foreshortening would have been a predicted convergent adaptation to harder bites, as it evolves frequently in response to selection for increased mechanical advantage (Goswami et al., 2010; Mitchell et al., 2018; Mitchell, Sherratt & Weisbecker, 2024d; Wroe & Milne, 2007). Such “many-to-one mapping” cases of multiple morphologies performing similar functions will be invisible to assessments of morphological convergence, which focus on shape similarities between taxa weighted by phylogenetic distance (Chatar, Fischer & Tseng, 2022; Grossnickle et al., 2024; Law, Linden & Flynn, 2024; Mitchell et al., 2024a; Stayton, 2015).

It is possible that the lack of facial foreshortening in *B. penicillata* might relate to its dependence on subterranean fungal sporocarps (i.e. truffles). Bettongs and potoroos most likely detect ripe sporocarps with olfactory cues (Donaldson & Stoddart, 1994; Vernes & Jarman, 2014). Shortening of the face might reduce the surface area of turbinates committed to olfactory sensation (Van Valkenburgh, Pang, Bird, Curtis, Yee, Wysocki & Craven, 2014), which may have constrained the facial shape of *B. penicillata. Bettongia lesueur* relies on truffles substantially less than the other three species (∼5% of diet; Mitchell et al., 2024a) and might thus be less susceptible to selection for high-performance olfaction, resulting in a morphology better suited for other functions. Although this study did not investigate the relative size of the turbinates (olfactory vs. respiratory) in bettongs, this could offer a useful avenue for further research to understand potentially competing demands involved with foraging behaviour of bettongs.

Considering most of the mechanical effort to crack open *Santalum* seeds is performed at the P3, the facial proportions anterior to this region are not directly involved in *Santalum* cracking and might thus reflect adaptations to contrasting feeding ecology. While this might permit a longer rostrum for better olfaction in *B. penicillata*, the facial skeleton of *B. lesueur* anterior to the premolars is more reminiscent of browsing morphology seen in kangaroos and other distant relatives of potoroids (Macropodiformes), and this aligns with a higher proportion of browse in this species’ diet compared to the other two species (Dawson & Milne, 2012; Mitchell et al., 2018). These results therefore have potential implications for the conservation of *B. penicillata*, which has been translocated into arid zone sites unlikely to have sites with an abundance in fungi required by this species to thrive. If strong selection for olfactory sensation of fungal sporocarps is confirmed for *B. penicillata*, evaluation of site fungal abundance would likely prove to be an important consideration before translocating the species (Bougher, 2008).

Finite Element Analyses (FEA) have shown that the species with the lowest mechanical efficiency in our sample is the Tasmanian bettong, *B. gaimardi* (Mitchell et al., 2025), whose anteriorly placed dentition reduces its relative bite force compared to the shorter-faced species in the sample. This might be related to the advantages of longer faces in obtaining food in the absence of selection for harder bites (Mitchell, Sherratt & Weisbecker, 2024c).

Importantly, Tasmanian bettongs also have the largest crania in our sample, which means they also likely have naturally overall larger muscles. Because bite force is a product of muscle force and mechanical advantage (Popowics & Herring, 2006), and muscle force depends on overall muscle size, this means that a species with a larger cranium and a longer face can still exert the same bite forces as a smaller species with a shorter face (Mitchell et al., 2024c). A clustering of *B. gaimardi* in PC2 maximum, representing more anteriorly extended P3 premolars, might therefore reflect some lower bite force demands at the premolar region, owing to its larger cranial sizes (Mitchell et al., 2024b).

A final intriguing observation is a striking difference in shape between *B. tropica* specimens that were from the wild, compared to specimens that were from a captive colony. Despite the small sample size, captives clearly occupied a unique region of morphospace between *B. lesueur* and the other two species, whereas wild specimens were mostly intermediate in shape to *B. penicillata* and *B. gaimardi.* The captive animals were born in captivity to two wild caught individuals captured in Queensland (Smith, 1998). Eleven captive-born *B. tropica* were fostered with a *B. penicillata* foster mother, while another six were weaned by their wild caught mother (Smith, 1998); however, we were unable to ascertain which specimens were cross-fostered. Their adult diet consisted of apples, bananas, boiled potato, dry dog food, and mixed large seeds, especially sunflower seeds (Smith, 1998). It is likely that either the cross-fostering or the mechanical properties of the captive dietary regime led to the cranial shape differences of captive animals relative to the wild ones. Dietary differences within species have also been suggested for differences in cranial shape in other marsupials (Mitchell et al., 2020; Weisbecker et al., 2019) and many other mammals (e.g., Curtis et al., 2018; Hartstone-Rose et al., 2014; Mitchell et al, 2021). As can be seen in our study, these differences can be extensive and represent an issue for captive breeding but also have substantial potential for confounding the study of extinct mammals (Mitchell et al., 2024a).

Importantly, this effect also suggests substantial capacity of individuals within a species to prevail changing dietary conditions. It is possible that developmental responses, such as increases in bone deposition in areas under high strain, support a wide range of feeding behaviours, for example by compensating for cranial shapes with lower resilience to strain as suggested for *B. penicillata* above. This has important implications for the planning of dietary regimes in conservation efforts (e.g., Siciliano-Martina, Light & Lawing, 2021). It is also relevant for taxonomic assessments of mammalian crania, which require consideration of phenotypic plasticity as a potential source non-informative, within-taxon variation (Viacava, Baker, Blomberg, Phillips & Weisbecker, 2022), and also palaeoecological assessments of extinct species.

## CONCLUSION

We identified clear morphological adaptations to generating high bite forces in the two species of bettongs that crack open *Santalum* seeds but unexpectedly found that these adaptations were not related to morphological convergence. This unexpected case of many-to-one mapping of cranial shape to diet, at a fine taxonomic scale, calls for nuanced considerations of relating feeding ecology to skull morphology in mammals. In the case of these bettongs, the partial overlap of diets in *B. penicillata* and *B. lesueur* would have almost certainly gone undetected under the assumptions of previously ascribed diet categorisations. A further complicating factor is the clear developmental susceptibility of cranial shape to captivity, suggesting that the same species can display substantial differences in cranial shape according to environments. Further assessments of olfactory sensation in *Bettongia* spp. might reveal the importance of fungal diversity in determining site adequacy for translocations.

## DATA AVAILABILITY STATEMENT

All scans will be made available on Morphosource following publication. All data and analyses are available on GitHub: https://github.com/Maddison0137/Bettong_skull_shape_study

## REFERENCES

1. Adams DC, Otárola-Castillo E, Paradis E. 2013. Geomorph: an R package for the collection and analysis of geometric morphometric shape data. Methods in Ecology and Evolution 4: 393–399.

2. Alexander RM. 1981. Factors of safety in the structure of animals. Science Progress 67: 109–130.

3. Archer M. 1979. *Wabularoo naughtoni* gen. et sp. nov., an enigmatic kangaroo (Marsupialia) from the middle Tertiary Carl Creek Limestone of northwestern Queensland. Memoirs of the Queensland Museum 19: 299–307.

4. Arman SD, Prideaux GJ. 2015. Dietary classification of extant kangaroos and their relatives (Marsupialia: Macropodoidea). Austral Ecology 40: 909–922.

5. Bice J, Moseby K. 2008. Diets of the re-introduced greater bilby (Macrotis lagotis) and burrowing bettong (Bettongia lesueur) in the Arid Recovery Reserve, Northern South Australia. Australian Mammalogy 30.

6. Biknevicius AR, Ruff CB. 1992. The structure of the mandibular corpus and its relationship to feeding behaviours in extant carnivorans. Journal of Zoology 228: 479–507.

7. Bougher N L, Friend T, Bell L. 2008. Fungi available to and consumed by translocated Gilbert’s potoroos: preliminary assessments at three translocation sites. Department of Environment and Conservation, Western Australia. [online] Available at: https://library.dbca.wa.gov.au/FullTextFiles/024745.pdf [Accessed 18 July 2025]

8. Butler K, Travouillon KJ, Price GJ, Archer M, Hand SJ. 2016. Cookeroo, a new genus of fossil kangaroo (Marsupialia, Macropodidae) from the Oligo-Miocene of Riversleigh, northwestern Queensland, Australia. Journal of Vertebrate Paleontology 36.

9. Case JA. 1984. A new genus of Potoroinae (Marsupialia: Macropodidae) from the Miocene Ngapakaldi Local Fauna, South Australia, and a definition of the Potoroinae. Journal of Paleontology 58: 1074–1086.

10. Chapman A. (2025). Bettongia penicillata (Woylie). [online] Available at: https://www.flickr.com/photos/arthur_chapman/3009409238 [Accessed 17 Jul. 2025].

11. Chapman TF. 2015. Reintroduced burrowing bettongs (Bettongia lesueur) scatter hoard sandalwood (Santalum spicatum) seed. Australian Journal of Zoology 63.

12. Chatar N, Fischer V, Tseng ZJ 2022. Many-to-one function of cat-like mandibles highlights a continuum of sabre-tooth adaptations. Proceedings of the Royal Society B 289: 20221627.

13. Collyer ML, Adams DC. 2018. RRPP: An r package for fitting linear models to high-dimensional data using residual randomization. Methods in Ecology and Evolution 9: 1772–1779.

14. Constantino PJ, Wright BW. 2009. The importance of fallback foods in primate ecology and evolution. Am J Phys Anthropol 140: 599–602.

15. Curtis AA, Orke M, Tetradis S, van Valkenburgh B. 2018. Diet-related differences in craniodental morphology between captive-reared and wild coyotes, Canis latrans (Carnivora: Canidae). Biological Journal of the Linnean Society 123: 677–693.

16. Dawson R, Milne N. 2012. Cranial size and shape variation in mainland and island populations of the quokka. Journal of Zoology 288: 267–274.

17. Donaldson R, Stoddart M. 1994. Detection of hypogeous fungi by Tasmanian bettong (Bettongia gaimardi: Marsupialia; Macropodoidea). Journal of Chemical Ecology 20: 1201–1207.

18. Figueirido B, Serrano-Alarcón FJ, Palmqvist P, Kitchener A. 2011. Geometric morphometrics shows differences and similarities in skull shape between the red and giant pandas. Journal of Zoology 286: 293–302.

19. Figueirido B, Tseng ZJ, Martin-Serra A. 2013. Skull shape evolution in durophagous carnivorans. Evolution 67: 1975–1993.

20. Flannery TF, Archer M. 1987. *Bettongia moyesi*, a new and plesiomorphic kangaroo (Marsupialia: Potoroidae) from Miocene sediments of northwestern Queensland. In: Archer M, ed. Possums and Opossums: Studies in Evolution. Sydney: Surrey Beatty & Sons.

21. Flannery TF, Archer M, Plane M. 1983. Middle Miocene kangaroos (Macropodoidea: Marsupialia) from three localities in northern Australia, with a description nof two new subfamilies. Bureau of Mineral Resources Journal of Australian Geology and Geophysics 7: 287–302.

22. Frost HM. 1994. Wolff’s law and bone’s structural adaptations to mechanical usage: an overview for clinicians. The Angle Orthodontist 64: 175–188.

23. Goswami A, Milne N, Wroe S. 2010. Biting through constraints: cranial morphology, disparity and convergence across living and fossil carnivorous mammals. Proc Biol Sci 278: 1831–1839.

24. Grossnickle DM, Brightly WH, Weaver LN, Stanchak KE, Roston RA, Pevsner SK, Stayton CT, Polly PD, and Law CJ. 2024. Challenges and advances in measuring phenotypic convergence. Evolution 78: 1355–1371.

25. Haouchar D, Pacioni C, Haile J, McDowell MC, Baynes A, Phillips MJ, Austin JJ, Pope LC, Bunce M. 2016. Ancient DNA reveals complexity in the evolutionary history and taxonomy of the endangered Australian brush-tailed bettongs (Bettongia: Marsupialia: Macropodidae: Potoroinae). Biodiversity and Conservation 25: 2907–2927.

26. Harrison JJ. 2010. File:Bettongia gaimardi.jpg - Wikimedia Commons. [online] Available at: https://commons.wikimedia.org/wiki/File:Bettongia_gaimardi.jpg [Accessed 17 Jul. 2025].

27. Hartstone-Rose A, Selvey H, Villari JR, Atwell M, Schmidt T. 2014. The three-dimensional morphological effects of captivity. PLoS One 9: e113437.

28. Hunt, T. (2025). bettonger og potorooer – Lex. [online] Available at: https://lex.dk/bettonger_og_potorooer [Accessed 23 Jul. 2025].

29. Hylander WL, Johnson KR. 1997. In Vivo Bone Strain Patterns in the Zygomatic Arch of Macaques and the Significance of These Patterns for Functional Interpretations of Craniofacial Form. American journal of physical anthropology 102: 203–232.

30. Janis CM. 1990. Correlation of cranial and dental variables with dietary preferences in mammals: a comparison of macropodoids and ungulates. Memoirs of the Queensland Museum 28: 349–366.

31. Kaitlyn. 2010. File:Bettongia tropica 73525933.jpg - Wikimedia Commons. [online] Available at: https://commons.wikimedia.org/wiki/File:Bettongia_tropica_73525933.jpg [Accessed 17 Jul. 2025].

32. Kear BP, Pledge NS. 2007. A new fossil kangaroo from the Oligocene-Miocene Etadunna formation of Ngama Quarry, Lake Palankarinna, South Australia. Australian Journal of Zoology 55: 331–339.

33. Klaczko J, Sherratt E, Setz EZ. 2016. Are diet preferences associated to skulls shape diversification in xenodontine snakes? PLoS One 11: e0148375.

34. Klingenberg CP. 2016. Size, shape, and form: concepts of allometry in geometric morphometrics. Dev Genes Evol 226: 113–137.

35. Klingenberg CP, Barluenga M, Meyer A. 2002. Shape analysis of symmetric structures: quantifying variation among individuals and asymmetry. Evolution 56: 1909–1920.

36. Law CJ, Linden TJ, Flynn JJ. 2024. Skull evolution and lineage diversification in endemic Malagasy carnivorans. Royal Society Open Science 11: p.240538.

37. Mardia KV, Bookstein F, Moreton IJ. 2000. Statistical assessment of bilateral symmetry of shapes. Biometrika 87: 285–300.

38. McDowell MC, Haouchar D, Aplin KP, Bunce M, Baynes A, Prideaux GJ. 2015. Morphological and molecular evidence supports specific recognition of the recently extinctBettongia anhydra(Marsupialia: Macropodidae). Journal of Mammalogy 96: 287–296.

39. McIlwee AP, Johnson CN. 2002. The contribution of fungus to the diets of three mycophagous marsupials in Eucalyptus forests, revealed by stable isotope analysis. Functional Ecology 12: 223–231.

40. McNamara JA. 2014. Bettong diet and dentition. South Australian Naturalist, The 88: 80–90.

41. Mein E, Manne T, Veth P, Weisbecker V. 2022. Morphometric classification of kangaroo bones reveals paleoecological change in northwest Australia during the terminal Pleistocene. Scientific Reports 12: 18245.

42. Mitchell DR. 2019. The anatomy of a crushing bite: The specialised cranial mechanics of a giant extinct kangaroo. PLoS One 14: e0221287.

43. Mitchell DR, Kirchhoff CA, Cooke SB, Terhune CE. 2021. Bolstering geometric morphometrics sample sizes with damaged and pathologic specimens: Is near enough good enough? J Anat 238: 1444–1455.

44. Mitchell DR, Ledogar JA, Andrew D, Mathewson I, Weisbecker V, Vernes K. 2024a. The mechanical properties of bettong and potoroo foods. Australian Mammalogy 46.

45. Mitchell DR, Potter S, Eldridge MDB, Martin M, Weisbecker V. 2024b. Functionally mediated cranial allometry evidenced in a genus of rock-wallabies. Biology Letters 20: 20240045.

46. Mitchell DR, Sherratt E, Ledogar JA, Wroe S. 2018. The biomechanics of foraging determines face length among kangaroos and their relatives. Proc Biol Sci 285.

47. Mitchell DR, Sherratt E, Sansalone G, Ledogar JA, Flavel RJ, Wroe S. 2020. Feeding Biomechanics Influences Craniofacial Morphology at the Subspecies Scale among Australian Pademelons (Macropodidae: Thylogale). Journal of Mammalian Evolution 27: 199–209.

48. Mitchell DR, Sherratt E, Weisbecker V. 2024c. Facing the facts: adaptive trade-offs along body size ranges determine mammalian craniofacial scaling. Biol Rev Camb Philos Soc 99: 496–524.

49. Mitchell DR, Sherratt E, Weisbecker V. 2024d. Facing the facts: Adaptive trade-offs along body size ranges determine mammalian craniofacial scaling. Biological Reviews 99: 496–524.

50. Mitchell DR, Wroe S, Martin M, Weisbecker V. 2025. Testing hypotheses of skull function with comparative finite element analysis: three methods reveal contrasting results. Journal of Experimental Biology: jeb. 249747.

51. Mitchell DR, Wroe S, Ravosa MJ, Menegaz RA. 2021. More Challenging Diets Sustain Feeding Performance: Applications Toward the Captive Rearing of Wildlife. Integrative Organismal Biology 3: obab030.

52. Murphy M, Howard K, Hardy GESJ, Dell B. 2015. When losing your nuts increases your reproductive success: sandalwood (Santalum spicatum) nut caching by the woylie (Bettongia penicillata). Pacific Conservation Biology 21: 243–252.

53. Murphy MT, Garkaklis MJ, Hardy GESJ. 2005. Seed caching by woylies *Bettongia penicillata* can increase sandalwood *Santalum spicatum* regeneration in Western Australia. Austral Ecology 30: 747–755.

54. Page KD, Ruykys L, Miller DW, Adams PJ, Bateman PW, Fleming PA. 2019. Influences of behaviour and physiology on body mass gain in the woylie (Bettongia penicillata ogilbyi) post-translocation. Wildlife Research 46: 429–443.

55. Popowics T, Herring S. 2006. Teeth, jaws and muscles in mammalian mastication Feeding in domestic vertebrates: from structure to behaviour: CABI Wallingford UK. 61–83.

56. Prideaux GJ. 1999. *Borungaboodie hatcheri* gen. et sp. nov., a very large bettong (Marsupialia: Macropodoidea) from the Pleistocene of southwestern Australia. Records of the Western Australian Museum Supplement 57: 317–329.

57. R Development Core Team. 2023. R: A language and environment for statistical computing. Vienna, Austria: R Foundation for Statistical Computing.

58. Rick K, Ottewell K, Lohr C, Thavornkanlapachai R, Byrne M, Kennington WJ. 2019. Population Genomics of Bettongia lesueur: Admixing Increases Genetic Diversity with no Evidence of Outbreeding Depression. Genes (Basel) 10.

59. Robinson BW, Wilson DS. 1998. Optimal foraging, specialization, and a solution to Liem’s paradox. Am Nat 151: 223–235.

60. Robley AJ, Short J, Bradley S. 2008. Dietary overlap between the burrowing bettong (*Bettongia lesueur*) and the European rabbit (*Oryctolagus cuniculus*) in semi-arid coastal Western Australia. Wildlife Research 28: 341–349.

61. Rohlf FJ, Slice D. 1990. Extensions of the Procrustes Method for the Optimal Superimposition of Landmarks. Systematic Zoology 39: 40.

62. Rose RW, Rose RK. 1998. Bettongia gaimardi. Mammalian Species: 1–6.

63. Samuels JX. 2009. Cranial morphology and dietary habits of rodents. Zoological Journal of the Linnean Society 156: 864–888.

64. Sansalone G, Wroe S, Coates G, Attard MRG, Fruciano C. 2024. Unexpectedly uneven distribution of functional trade-offs explains cranial morphological diversity in carnivores. Nat Commun 15: 3275.

65. Sanson G. 1989. Morphological adaptations of teeth to diets in macropods. In: Grigg G, Jarman PJ and Hume ID, eds. Kangaroos, wallabies and rat-kangaroos. NSW, Australia: Surrey Beatty & Sons Pty Ltd. 151–168.

66. Santana SE, Dumont ER, Davis JL. 2010. Mechanics of bite force production and its relationship to diet in bats. Functional Ecology 24: 776–784.

67. Santana SE, Grosse IR, Dumont ER. 2012. Dietary hardness, loading behavior, and the evolution of skull form in bats. Evolution 66: 2587–2598.

68. Schlager S, Jefferis G, Ian D. 2018. Package ‘Morpho’.

69. Siciliano-Martina L, Light JE, Lawing AM. 2021. Changes in canid cranial morphology induced by captivity and conservation implications. Biological Conservation 257.

70. Smith MJ. 1998. Establishment of a captive colony of Bettongia tropica (Marsupialia: Potoroidae) by cross-fostering; and observations on reproduction. Journal of Zoology 244: 43–50.

71. Stayton CT. 2015. The definition, recognition, and interpretation of convergent evolution, and two new measures for quantifying and assessing the significance of convergence. Evolution 69: 2140–2153.

72. Taylor RJ. 1992. Seasonal Changes in the Diet of the Tasmanian Bettong (*Bettongia gaimardi*), a Mycophagous Marsupial. Journal of Mammalogy 73: 408–414.

73. Ungar PS, Grine FE, Teaford MF. 2008. Dental microwear and diet of the Plio-Pleistocene hominin Paranthropus boisei. PLoS One 3: e2044.

74. Van Valkenburgh B, Pang B, Bird D, Curtis A, Yee K, Wysocki C, Craven BA. 2014. Respiratory and olfactory turbinals in feliform and caniform carnivorans: the influence of snout length. The Anatomical Record 297: 2065–2079.

75. Vernes K, Castellano M, Johnson CN. 2002. Effects of season and fire on the diversity of hypogeous fungi consumed by a tropical mycophagous marsupial. Journal of Animal Ecology 70: 945–954.

76. Vernes K, Jarman P. 2014. Long-nosed potoroo (Potorous tridactylus) behaviour and handling times when foraging for buried truffles. Australian Mammalogy 36.

77. Viacava P, Baker AM, Blomberg SP, Phillips MJ, Weisbecker V. 2022. Using 3D geometric morphometrics to aid taxonomic and ecological understanding of a recent speciation event within a small Australian marsupial (Antechinus: Dasyuridae). Zoological Journal of the Linnean Society 196: 963–978.

78. Viacava P, Blomberg SP, Sansalone G, Phillips MJ, Guillerme T, Cameron SF, Wilson RS, Weisbecker V. 2020. Skull shape of a widely distributed, endangered marsupial reveals little evidence of local adaptation between fragmented populations. Ecol Evol 10: 9707–9720.

79. Weisbecker V, Guillerme T, Speck C, Sherratt E, Abraha HM, Sharp AC, Terhune CE, Collins S, Johnston S, Panagiotopoulou O. 2019. Individual variation of the masticatory system dominates 3D skull shape in the herbivory-adapted marsupial wombats. Front Zool 16: 41.

80. Westerman M, Loke S, Tan MH, Kear BP. 2022. Mitogenome of the extinct Desert ‘rat-kangaroo’ times the adaptation to aridity in macropodoids. Sci Rep 12: 5829.

81. Willie BM, Zimmermann EA, Vitienes I, Main RP, Komarova SV. 2020. Bone adaptation: Safety factors and load predictability in shaping skeletal form. Bone 131: 115114.

82. Woodburne MO. 1984. *Wakiewakie lawsoni*, a new genus and species of Potoroinae (Marsupialia: Macropodidae) of medial Miocene age, South Australia. Journal of Paleontology 58: 1062–1073.

83. Wroe S, Milne N. 2007. Convergence and remarkably consistent constraint in the evolution of carnivore skull shape. Evolution 61: 1251–1260.

84. Zelditch ML, Ye J, Mitchell JS, Swiderski DL. 2017. Rare ecomorphological convergence on a complex adaptive landscape: Body size and diet mediate evolution of jaw shape in squirrels (Sciuridae). Evolution 71: 633–649.

85. Zosky KL, Wayne AF, Bryant KA, Calver MC, Scarff FR. 2017. Diet of the critically endangered woylie (Bettongia penicillata ogilbyi) in south-western Australia. Australian Journal of Zoology 65.

